# GOBoost: Leveraging Long-Tail Gene Ontology Terms for Accurate Protein Function Prediction

**DOI:** 10.1101/2024.11.16.623961

**Authors:** Lei Zhang, Yang Wang, Xiao Chen, Jie Hou, Dong Si, Rui Ding, Bo Jiang, Hailey Ledenko, Renzhi Cao

## Abstract

**Motivation:** With the advancement of deep learning, researchers have increasingly proposed computational methods based on deep learning techniques to predict protein function. However, many of these methods treat protein function prediction as a multi-label classification problem, often overlooking the long-tail distribution of functional labels (i.e., Gene Ontology Terms) in datasets. To address this issue, we propose the GOBoost method, which incorporates the proposed long-tail optimization ensemble strategy. Besides, GOBoost introduces the proposed global-local label graph module and multi-granularity focal loss function to enhance long-tail functional information, mitigate the long-tail phenomenon, and improve overall prediction accuracy.

**Results:** We evaluate GOBoost and other state-of-the-art (SOTA) protein function prediction methods on the PDB and AF2 datasets. The GOBoost outperformed SOTA methods across all evaluation metrics on both datasets. Notably, in the AUPR evaluation on the PDB test set, GOBoost improved by 10.71%, 35.91%, and 22.71% compared to the SOTA HEAL method in the MF, BP, and CC functions. The experimental results demonstrate the necessity and superiority of designing models from the label long-tail distribution perspective.

**Availability:** https://github.com/Cao-Labs/GOBoost

## Introduction

Proteins are essential molecules in biological processes, playing critical roles in catalysis, recognition, and transmission of signals, and providing structural support for cells and tissues [1]. Understanding protein function is highly significant for biomedical applications. Many diseases are associated with abnormal protein function, including diabetes and Parkinson’s disease [2]. By advancing our knowledge of protein function, we can uncover disease mechanisms and facilitate drug development. The Gene Ontology [3] Consortium has established a standardized functional vocabulary library, Gene Ontology, to define and describe specific functions. These functions are categorized into three sub-functional ontologies: molecular function (MF), biological process (BP), and cellular component (CC). With advancements in protein sequencing technology [4, 5], the number of protein sequences has increased dramatically. Although the traditional experimental method of annotating protein functions is accurate, it is time-consuming and labor-intensive. Therefore, there is an increasing gap between the number of surging protein sequences and the number of proteins that have been experimentally annotated [6]. Therefore, it is crucial to develop a fast and accurate computational method to classify protein’s function type.

Knowledge transfer-based functional prediction methods are usually based on the assumption that proteins with high sequence similarity tend to exhibit similar functional properties. BLAST [7] and DIAMOND [8] perform sequence comparisons by querying proteins and their known functions in a database, using sequence similarity as the transition probability for functional annotation. FunFam [9] transfers known functions by identifying similar structural domains in the database. These methods depend on prior database knowledge, and when faced with large-scale data processing, the process of matching the database can be highly time-consuming. With advancements in deep learning technology, researchers are increasingly focusing on developing computational methods that address these issues. Protein sequences are the most commonly used feature information for protein function prediction. CNN-based [10] methods like DeepGO [11], DeepGOPlus [12], and SDN2GO [13] capture functional motifs of different lengths in protein sequences through convolution operations, establishing mapping relationships between these motifs and corresponding functions. Other methods [14, 15] use RNN-like [16] models to more effectively capture contextual information in sequence data. Transformer [17] technology has gained popularity due to its attention mechanism, which can capture relationships between long-range residues in protein sequences. TALE [18] uses a transformer encoder to capture long-range relationships between amino acids. The latest research [19, 20, 21, 22] shows that using the pre-trained biological large language model [23, 24, 25] based on the transformer architecture to extract sequence features and applying these features to downstream function prediction tasks can achieve better results.

The above sequence-based methods are easily affected by amino acid variations, while the 3D structure is often more conserved than the sequence and is usually closely related to protein function. To this end, GAT-GO [26] uses RaptorX [27] to directly predict the amino acid contact graph in the three-dimensional structure of proteins. With the development of high-quality protein structure prediction methods [28], researchers generally use the 3D structure of proteins to more accurately abstract amino acid contact graphs. On this basis, various graph convolutional networks [26, 29] are widely used to mine the complex relationship between protein structure information and function. Therefore, this type of methods [30, 26, 31, 32, 21] can generally be summarized as a graph multi-label classification problem. Considering the natural characteristics of multi-label classification problems, some researchers have tried to use the information between labels to improve the prediction performance of the model. One approach, known as the hierarchical output methods [11, 33], directly utilizes the hierarchical constraint information of the functional Directed Acyclic Graph (DAG) to update prediction probabilities, thereby avoiding sub-optimal solutions. Another approach does not strictly incorporate the hierarchical constraint information of the functional DAG. They usually update the functional label information by analyzing the associations between various labels. PANDA2 [20] and PANDA-3D [34] use the bidirectional DAG of GO terms and the GO terms relationship implicitly captured by the transformer decoder to update the label information, respectively. DeepFPF-CO [35] uses the prior knowledge of GO terms from the training set to construct a conditional co-occurrence matrix to update the predicted probability. At the same time, DeepFPF-CO also effectively reduces the noise introduced by low-frequency terms by setting a specific threshold.

The computational methods discussed above often treat the protein function prediction problem as a straightforward multi-label classification task. Due to how the functional DAG is constructed, functions at more profound levels are usually more specific. However, only a few proteins are annotated with these specific functions. Consequently, this leads to a pronounced long-tail phenomenon in the overall distribution of GO terms within the dataset, as illustrated in Fig. 1. Few terms are highly frequent, while the majority appear infrequently. Therefore, adopting a simple multi-label classification approach may cause the model to ignore these more specific low-frequency functions. To address this, Al-Shahab et al. [36] and Jaehee Jung et al. [37] proposed the under-sampling technique to balance the distribution of GO terms. However, with the continuous expansion in the number of functional categories and expansion of the dataset, it is becoming increasingly difficult to mitigate the long-tail phenomenon effect using sampling technology.

**Fig. 1.**
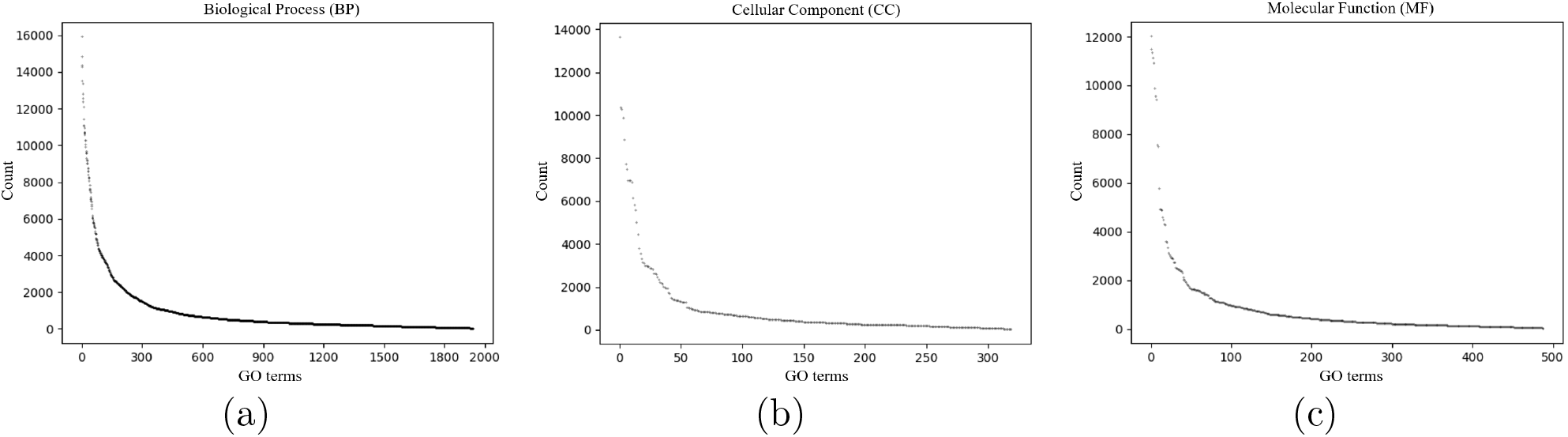
The dots is GO terms; the y-axis is the number of times the GO terms appear in the PDB and AF2 training sets, and the x-axis is the order of the terms sorted from largest to smallest according to the number of occurrences. **(a)** Statistics on BP GO terms. **(b)** Statistics on CC GO terms. **(c)** Statistics on MF GO terms.

To address the above problems, we proposed GOBoost, a novel protein function prediction method designed to alleviate the long-tail effect. This method introduces an optimized ensemble strategy tailored to the long-tail distribution, integrating three complementary base models: GOBoost^Head^, which is trained on high-frequency labels; GOBoost^Tail^, which is trained on medium and low-frequency labels; and GOBoost^All^, which is trained on all labels. The base model utilizes Class Activation Mapping (CAM) [38] technology to map protein structure features encoded by a Graph Convolutional Network [29] into functional label features. CAM uses a classifier to identify class-specific activation scores, which are then used to weight the protein features, resulting in content-aware label features. These features are passed through the proposed global label graph module to capture the co-occurrence relationships among global high-frequency functional labels, while the proposed local individual functional label graph is constructed to supplement relationships involving low-frequency labels. A multi-granularity focal loss function is used during training, assigning a greater weight to long-tail labels, thereby encouraging the model to focus on these underrepresented labels. Experimental results demonstrated that GOBoost, specifically designed to address long-tail distribution, achieved higher precision accuracy and superior generalization compared to other state-of-the-art methods.

## Materials and methods

### Datasets

This method uses the same Protein Data Bank (PDB) dataset [30] as DeepFRI and AlphaFold 2 (AF2) dataset (same as what HEAL used) [21]. The PDB dataset consists of natural protein structures obtained through physical experiments, which are divided into training, validation, and testing sets in a sequential 8:1:1 ratio. All annotated protein function identifiers are sourced from SIFTS (2019-06-18) [39], which transfers functions to PDB chains according to residue mappings between UniProtKB (release 2019-06) [40] and PDB (release 24.19 version) entries. The training and validation sets use both EXP and IEA GO annotations, while the test set relies solely on EXP annotations. Additionally, each protein in the PDB contains at least one experimentally determined GO term across each functional ontology. HEAL uses homologous proteins of PDB as the enhanced dataset AF2. The protein structures in this dataset were predicted by AlphaFold 2, and each protein was annotated with at least one GO term with an IC (Information Content [41]) *>* 10. Subsequently, MMseqs [42] was used to cluster protein sequences with a 25% sequence similarity, resulting in 38,185 training proteins, 4,242 validation proteins, and 567 test proteins.

To improve the dynamic learning of the co-occurrence relationship between labels, we further processed the PDB and AF2 datasets. All training and validation sets were preprocessed according to three sub-functional ontologies by removing all proteins without corresponding functions (i.e., all proteins without positive labels were excluded). To ensure a fair evaluation, the test sets of PDB and AF2 datasets are consistent with DeepFRI and HEAL. Unlike the PDB test set, the AF2 test set may contain proteins without corresponding subfunctional labels, with 137, 9, and 43 proteins lacking corresponding MF, BP, and CC functions, respectively. Detailed data statistics are shown in Table 1.

**Table 1.** Protein data statistics on PDB and AF2 datasets.

| DataSet | Statistic | MF | BP | CC | All |
| --- | --- | --- | --- | --- | --- |
| BP | Training Size | 24,956(83.48%) | 23,513(78.66%) | 11,295(37.78%) | 29,893 |
|  | Validating Size | 2,749(82.75%) | 2,624(78.99%) | 1,299(39.10%) | 3,322 |
|  | Testing Size | 3,414(100%) | 3,414(100%) | 3,414(100%) | 3,414 |
|  | Number of classes | 489(17.77%) | 1,943(70.60%) | 320(11.63%) | 2,752 |
| AF2 | Training Size | 33,827(88.59%) | 37,263(97.59%) | 32,397(84.84%) | 38,185 |
|  | Validating Size | 3,708(87.41%) | 4,146(97.74%) | 3,663(86.35%) | 4,242 |
|  | Testing Size | 567(100%) | 567(100%) | 567(100%) | 567 |
|  | Number of classes | 489(17.77%) | 1,943(70.60%) | 320(11.63%) | 2,752 |

Note: This table 1 shows the number of proteins in the dataset and the number of their three types of functional labels, and records their proportion in the total.

### Model architecture

#### Overview

The framework of GOBoost is illustrated in Fig. 2, which primarily comprises two components: the ensemble and base models. The ensemble model integrates the prediction results of three base models, and each focuses on training on different labels: Head, Tail, and All. The base model first inputs protein sequences into a pre-trained protein large language model to extract embedding matrices. It then uses AlphaFold 2 [28] to predict 3D structures, which are abstracted as protein graphs. Subsequently, the protein graph and the further processed sequence features are embedded into a graph convolution network to derive the final graph embedding features. Then, CAM [38] is used to translate the graph embedding features into GO term label embeddings. The embeddings are then fed into the global-local label graph module, which learns the co-occurrence relationships among the GO terms. Finally, the prediction results derived from the label embeddings are linearly combined with those from the protein graph embeddings to generate the final prediction probability. This probability is used to calculate the ℒ_*mgfl*_ loss for training the model.

**Fig. 2.**
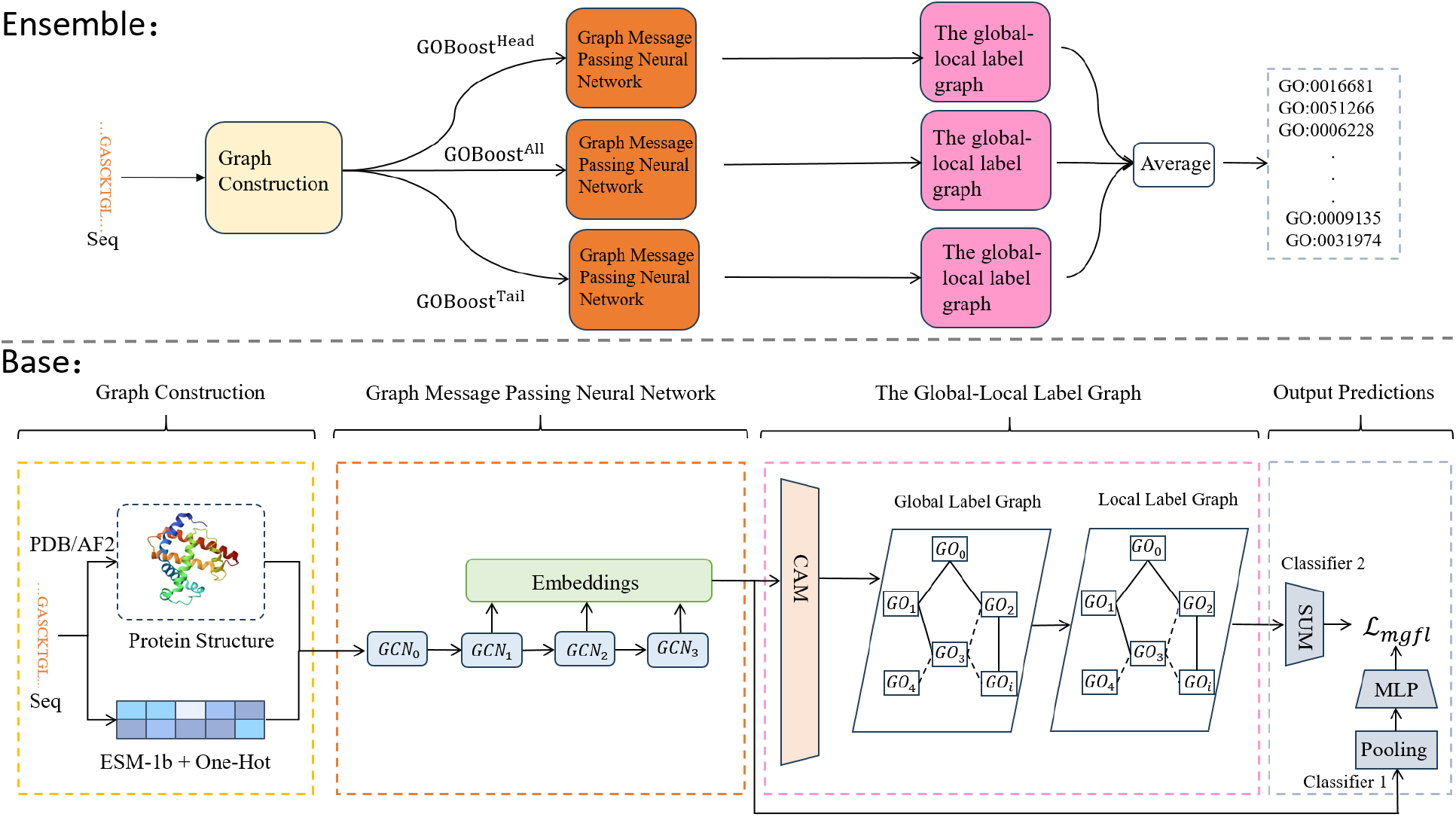
Overview of the proposed GOBoost. Ensemble: GOBoost is integrated from GOBoost^All^, GOBoost^Head^, and GOBoost^Tail^. Base: The base model learns the structural features of proteins and the co-occurrence relationship between GO terms to predict protein function.

#### Ensemble strategy

Due to the uneven distribution of GO terms, training a single model tends to overlook low-frequency GO terms. To address this, we proposed a long-tail optimization ensemble strategy that combines two base models, GOBoost^Head^ and GOBoost^Tail^, with the base model GOBoost^All^. All base models share the same model architecture and training loss function but are trained on different numbers of labels. GOBoost^All^ thoroughly captures relationships between all GO terms, while GOBoost^Head^ and GOBoost^Tail^ specialize in high-frequency GO terms and mid-to-low frequency GO term labels, respectively.

The ensemble strategy averages the overlapping prediction results from the three base models to produce the final prediction. GOBoost^Head^ and GOBoost^Tail^ effectively reduce the prediction error of GOBoost^All^, while GOBoost^All^ balances the predictions from GOBoost^Head^ and GOBoost^Tail^, thereby improving the robustness of the overall model, GOBoost. The probability of GO_*i*_ prediction after optimization integration is expressed as Formula 1:

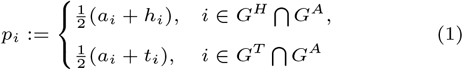

where *a*_*i*_, *h*_*i*_, and *t*_*i*_ denote the prediction values of GO_*i*_ given by All, Head, and Tail base models. *G*^*H*^ is the set of high-frequency GO term labels located in the “Head” region, *G*^*T*^ is the set of medium and low-frequency GO term labels located in the “Tail” region, and *G*^*A*^ is the set of all GO term labels.

#### Input features and graph message passing

### Residue embedding

In this work, we utilized the pre-trained protein large language model ESM-1b [24] to extract sequence-level features. Due to the limitation of sequence length in ESM-1b, we cut off the first 1000 residues to represent the entire protein sequence. The final sequence is input into the pre-trained ESM-1b model, and the output is extracted from the 33rd encoding layer. To represent amino acid types, this work uses a 0-20 encoding scheme. Then, PyTorch’s Embedding function is applied to map the discrete input (amino acid types) to a 96-dimensional continuous vector space. Finally, two MLPs are used to unify the 1280-dimensional ESM-1b embedding feature with the 96-dimensional amino acid species feature into a 512-dimensional vector. Then, these two types of features were linearly added to obtain the final protein residue level embedding *X* ∈ ℝ^*r*×*c*^.

### Graph construction

We abstract the 3D structure of proteins into an undirected graph G=(V, E, *X*^0^) with self-loops, where *V* denotes the graph nodes, *E* denotes the edges connecting nodes, and *X*^0^ denotes the initial feature matrix of the graph. Each node V_*i*_ corresponds to an amino acid, represented by the C_*α*_ atom of the residue. An edge E_*ij*_ is formed between two nodes if the Euclidean distance between their C_*αi*_ and C_*αj*_ atoms is less than 10*Å* (represented by a 0,1 matrix *A* ∈ ℝ^*r*×*r*^ to indicate the connection relationship between residues in the protein), and 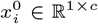 serves as the initial feature of node_*i*_.

### Graph message passing

The protein graph’s adjacency matrix *A* and initial feature *X* are fed into a graph convolution neural network, which assigns different aggregation weights to update the features of residue_*i*_ based on the degree information of adjacent residues. The receptive field range of convolution layer features varies across different layers. As the number of convolution layers increases, the receptive field range of deeper convolution features also expands. This work performs a weighted summation of the last L-1 layers of protein graph features through a learnable aggregation parameter to generate the final graph feature representation.

#### Global-local label graph

GOBoost uses CAM [38] to map residue-level protein features *H*^*f*^ ∈ ℝ^*r*×*c*^ to label features *H*^*g*^ ∈ ℝ^*g*×*c*^. Unlike the static co-occurrence matrix built based on prior knowledge [35], this module dynamically establishes the co-occurrence relationship between labels during training [43]. The label features are input into the global GO terms graph module. The label features are aggregated according to the global label co-occurrence matrix, followed by the parameter matrix to update the label features. The global label co-occurrence matrix is shared for each protein, so it tends to capture the co-occurrence relationship between high-frequency labels during training. However, many low-frequency GO terms still exist, and their co-occurrence relationships with other GO terms differ across proteins, making these relationships more complex for the model to capture.

Therefore, to address this, we established a local label graph. The input of the local label graph is formed by concatenating the label features extracted by the global label graph with its global features obtained by global average pooling. This module updates the label features using the label co-occurrence matrix and the parameter matrix. Unlike the global label co-occurrence matrix, the local individual label co-occurrence matrix is learned based on the characteristics of each protein through the parameter matrix, so each protein has a different local individual label co-occurrence matrix. This approach compensates for the low-frequency GO terms information that may be overlooked by the global label graph, effectively capturing the relationship between low-frequency GO terms and other GO terms, thus enhancing the overall predictive performance of the model.

#### Output predictions

### Graph pooling classification prediction (Classifier 1)

This process converts residue-level protein features into graph-level representations using average pooling operations, capturing more global information. This approach enables protein function prediction by analyzing overall trends in residue characteristics. After the graph pooling operation, the graph feature is input into the MLP layer and the linear activation function (RELU). A linear activation operation is performed after increasing the dimensionality of the graph features through the first layer of MLP. Finally, the graph features are input into the second layer of MLP and sigmoid function to obtain the predicted probabilities of all protein functions.

### Label feature classification prediction (Classifier 2)

The global-local label mapping module generates independent label features, enabling each to be directed into its respective classification head. During GO term probability prediction, each feature channel of the labeled features is linearly combined and then passed through a sigmoid function to yield the predicted probability of the protein function.

The final GO term predictions are generated by linearly combining the prediction probabilities from the two distinct classification heads, as shown in Formula 2:

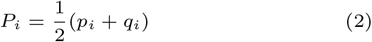

*P*_*i*_ is the predicted probability of *GO*_*i*_, *p*_*i*_ is the predicted probability of Classifier 1, and *q*_*i*_ is the predicted probability of Classifier 2.

#### Multi-grained focal loss function

The model aims to predict hundreds or even thousands of GO terms, though each protein generally corresponds to only a small subset of these terms. This leads to a severe imbalance between positive and negative labels during training. When calculating the loss, negative labels often dominate, which affects the model’s ability to predict positive labels. Additionally, many GO terms in the training set occur infrequently, leading the model to favor high-frequency terms during training while often overlooking these low-frequency yet more specific GO terms.

Specifically, the focal parameter *γ* in the traditional focal loss function [44] is refined into the positive and negative focal parameters *γ*^+(*i*)^ and *γ*^−(*i*)^ of each GO_*i*_. Then, parameters *γ*^+(*i*)^ and *γ*^−(*i*)^ are further decoupled into the above two granularities [45]: the positive and negative label focus parameter *γ*_*pn*_ and the head-tail focus parameter 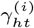. When *γ*^+(*i*)^ *< γ*^−(*i*)^, the contribution of negative labels in the loss can be reduced, allowing the model to focus on positive labels. Here, we set *γ*^+(*i*)^ = 0 to maximize the contribution of positive labels in the loss. For *γ*^−(*i*)^, the sum of 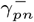 and 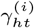 constitutes the focus parameter *γ*^−(*i*)^ of the negative label GO_*i*_. 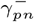 [46] is used to adjust the negative focus ratio of all negative labels, while 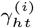 is responsible for adjusting the long-tail focus ratio of the negative label GO_*i*_. 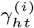 adjusts the head-tail label focus using the maximum regularization function [47] based on the training set’s static distribution of protein functions. A larger 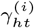 value for tail labels enhances the model’s sensitivity to these labels during gradient backpropagation, encouraging it to allocate more focus to long-tail labels. As shown in Formula 3, the loss function ℒ_*mgfl*_ is defined as follows:

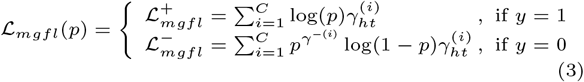

*C* is the number of functional categories, *i* represents the *i*^*th*^ protein function, *p* is the predicted probability, and *y* is the actual label (0, 1).

## Model training

GOBoost was initially trained on the PDB [30] dataset (i.e., GOBoost^All(PDB)^). Research shows that DeepFRI using homologous proteins for model training can significantly improve the prediction ability of the model. Based on this research result, we further integrated the AF2 data set containing PDB homologous proteins as enhanced training data, combined with the original PDB data set, to jointly train multiple model variants (i.e., GOBoost^All^, GOBoost^Head^, GOBoost^Tail^, and GOBoost). We trained all models on a host equipped with an NVIDIA RTX 3080 GPU (16GB video memory), with a batch size of 32 and a maximum training epoch of 100. An early stopping strategy with a window size of ten epochs is employed during training, where training is terminated if the validation loss does not decrease over any ten consecutive epochs. When training with only the PDB training set, we validate using only the PDB validation set. However, when both the PDB and AF2 training sets are used, the validation set includes both the PDB and AF2 validation sets. All base models of GOBoost are optimized using the Adam optimizer with a learning rate of 1e-4. According to the distribution trend of GO term occurrences in PDB and AF2 training sets in Fig. 1, GOBoost^Head^ roughly selects the top 100, 300, and 50 high-frequency labels for training on MF, BP, and CC functional ontologies, respectively. The remaining medium and low-frequency labels are used to train GOBoost^Tail^.

## Results and discussion

In order to comprehensively evaluate the performance of GOBoost, we first compared it with other advanced methods on the PDB and AF2 datasets. Then, we showed the specific advantages of GOBoost over the current SOTA HEAL method on the PDB dataset. Finally, we conducted a series of ablation experiments to analyze our designed module’s specific improvement effect in depth.

### Performance on the PDB dataset

As shown in Table 2, GOBoost achieved the best results across all evaluation metrics for the three sub-function ontologies. Specifically, GOBoost achieved scores of 0.765, 0.458, and 0.573 on AUPR, 0.787, 0.659, and 0.745 on Fmax, and 0.292, 0.450, and 0.401 on Smin for MF, BP, and CC functions, respectively. The prediction performance of GOBoost significantly surpassed Blast, FunFam, and DeepGO. This result shows that relying solely on protein sequence information to predict complex protein functions is not enough. Then, compared with the latest protein structure-based function prediction method HEAL, GOBoost increased AUPR scores by 10.71%, 35.91%, and 22.71% on the three sub-function ontologies, increased Fmax scores by 5.35%, 10.76%, and 8.44%, and decreased Smin scores by 14.62%, 11.59%, and 12.45%. Especially in dealing with the more challenging BP function, its performance improvement is particularly significant. Notably, GOBoost has the most significant improvement in AUPR, especially for BP, where it outperformed the HEAL method by 35.91%. It highlights GOBoost’s robustness, indicating that it effectively recognizes positive labels across various thresholds, handles imbalanced label data, and is not easily affected by threshold selection.

**Table 2.** Performance on the PDB DataSet.

| Method | AUPR ( ↑ ) | | | Fmax ( ↑ ) | | | Smin ( ↓ ) | | |
| --- | --- | --- | --- | --- | --- | --- | --- | --- | --- |
|  | MF | BP | CC | MF | BP | CC | MF | BP | CC |
| GOBoost | **0.765** | **0.458** | **0.573** | **0.787** | **0.659** | **0.745** | **0.292** | **0.450** | **0.401** |
| GOBoost<sup>All</sup> | 0.733 | 0.415 | 0.546 | 0.769 | 0.637 | 0.739 | 0.309 | 0.471 | 0.408 |
| GOBoost<sup>All(PDB)</sup> | 0.629 | 0.311 | 0.405 | 0.702 | 0.574 | 0.680 | 0.373 | 0.517 | 0.465 |
| Blast | 0.136 | 0.067 | 0.097 | 0.038 | 0.336 | 0.448 | 0.632 | 0.651 | 0.628 |
| FunFams | 0.367 | 0.260 | 0.228 | 0.572 | 0.500 | 0.627 | 0.531 | 0.579 | 0.503 |
| DeepGO | 0.391 | 0.182 | 0.263 | 0.577 | 0.493 | 0.594 | 0.472 | 0.577 | 0.550 |
| DeepFRI | 0.495 | 0.261 | 0.274 | 0.625 | 0.540 | 0.613 | 0.437 | 0.543 | 0.527 |
| HEAL<sup>PDB</sup> | 0.571 | 0.259 | 0.342 | 0.691 | 0.565 | 0.655 | 0.401 | 0.540 | 0.501 |
| HEAL | 0.691 | 0.337 | 0.467 | 0.747 | 0.595 | 0.687 | 0.342 | 0.509 | 0.458 |

Note: Data in bold indicates the best result. GOBoost and GOBoost<sup>All</sup> are trained on PDB and AF2 training datasets. GOBoost<sup>All(PDB)</sup> is trained only on the PDB training set. GOBoost<sup>All</sup> and GOBoost<sup>All(PDB)</sup> are base models without ensemble strategy.

GOBoost^All^ achieved scores of 0.733, 0.415, and 0.546 on AUPR, 0.769, 0.637, and 0.739 on Fmax, and 0.309, 0.471, and 0.408 on Smin, for the MF, BP, and CC function categories. Compared to the HEAL method, GOBoost^All^ increased AUPR by 6.08%, 23.15%, and 16.92%, respectively across the three function types. Fmax increased by 2.95%, 7.06%, and 7.57%, and Smin decreased by 9.65%, 7.47%, and 10.92%. Although the performance of GOBoost^All^ was slightly lower than GOBoost, it still outperformed the HEAL method in all evaluation metrics. The experimental data indicates a consistent improvement trend between GOBoost^All^ and GOBoost, with both achieving the best improvement effect on AUPR. Importantly, even without applying the long-tail optimization ensemble strategy proposed in this work, the base model still effectively handles this imbalanced data.

This work further analyzed the performance of our base model GOBoost^All(PDB)^. GOBoost^All(PDB)^ achieved scores of 0.629, 0.311, and 0.405 on AUPR, 0.702, 0.574, and 0.680 on Fmax, and 0.373, 0.517, and 0.465 on Smin for MF, BP, and CC functions, respectively. GOBoost^All-PDB^ showed superior predictive performance compared to Blast, Funfams, DeepGO, and DeepFRI. Compared to DeepFRI, GOBoost^All(PDB)^ showed an AUPR improvement of 27.07%, 19.16% and 47.81% across MF, BP, and CC, respectively. Fmax increased by 12.32%, 6.30% and 10.93%, and Smin decreased by 14.65%, 4.76% and 11.76%. Compared with HEAL^PDB^, GOBoost^All(PDB)^ consistently delivers better results in all evaluation metrics. These results indicate that the GOBoost base model outperforms other methods even when trained on limited amounts of data.

### Performance on the AF2 dataset

In the AF2 test set, GOBoost’s prediction performance was evaluated under more challenging conditions, where the similarity between the protein sequences in the AF2 test and training sets was less than 25%. To evaluate GOBoost’s reliance on sequence similarity, we introduced the DeepGOPlus [12] method for comparative analysis. DeepGOPlus combines deep learning techniques with transfer annotation based on sequence similarity, which often outperforms traditional methods that rely solely on sequence similarity for annotation transfer.

As shown in Table 3, on the AF2 test set, GOBoost achieved the best results across all evaluation metrics for the three sub-functional ontologies. GOBoost achieved scores of 0.582, 0.246, and 0.318 on AUPR and 0.556, 0.497, and 0.643 on Fmax for MF, BP, and CC functions. Compared with DeepGOPlus, GOBoost increased AUPR by 25.70%, 21.18%, and 19.10% and Fmax by 23.56%, 15.58%, and 13.40%. In the difficult situation of transferring annotations without high-similarity sequences, GOBoost reduces its dependence on sequence similarity by leveraging structural information and GO terms co-occurrence information. Compared to the HEAL method, GOBoost increased AUPR for three types of functions by 15.64%, 23.00%, and 10.80% and Fmax by 13.24%, 4.63%, and 4.92%. The significant improvement in AUPR further demonstrates GOBoost’s ability to recognize positive labels efficiently in difficult situations, making its prediction results robust.

**Table 3.** Performance on the AF2 DataSet.

| Method | MF |  | BP |  | CC |  |
| --- | --- | --- | --- | --- | --- | --- |
| | AUPR ( ↑ ) | Fmax ( ↑ ) | AUPR ( ↑ ) | Fmax ( ↑ ) | AUPR ( ↑ ) | Fmax ( ↑ ) |
| GOBoost | **0.582** | **0.556** | **0.246** | **0.497** | **0.318** | **0.643** |
| GOBoost<sup>All</sup> | 0.540 | 0.526 | 0.220 | 0.480 | 0.313 | 0.621 |
| GOBoost<sup>All(PDB)</sup> | 0.366 | 0.418 | 0.121 | 0.397 | 0.186 | 0.521 |
| DeepFRI | 0.342 | 0.398 | 0.114 | 0.387 | 0.192 | 0.536 |
| HEAL<sup>PDB</sup> | 0.345 | 0.414 | 0.104 | 0.408 | 0.141 | 0.513 |
| DeepGOPlus | 0.463 | 0.450 | 0.203 | 0.430 | 0.267 | 0.567 |
| HEAL | 0.502 | 0.491 | 0.200 | 0.475 | 0.287 | 0.614 |

Note: Data in bold indicates the best result. GOBoost and GOBoost<sup>All</sup> are trained on PDB and AF2 training datasets. GOBoost<sup>All(PDB)</sup> is trained only on the PDB training set. GOBoost<sup>All</sup> and GOBoost<sup>All(PDB)</sup> are base models without ensemble strategy.

The performance comparison between the GOBoost^All^ and all comparison methods was further analyzed. GOBoost^All^ achieved 0.540, 0.220, and 0.313 on AUPR and 0.526, 0.480, and 0.313 on Fmax for MF, BP, and CC functions, respectively. GOBoost^All^ consistently outperformed all comparison methods across all evaluation metrics for the three sub-functional ontologies. This may be because the long-tail strategy we designed prompts the base model to pay more attention to rare functions, thereby improving the overall prediction performance of the model. GOBoost^All(PDB)^ shows that it’s prediction performance on MF and BP functions surpassed that of DeepFRI, even when DeepFRI was trained with enhanced data. Similar results have also been achieved on CC. GOBoost^All(PDB)^ outperformed HEAL^PDB^ in most evaluation metrics, and only slightly lower than HEAL^PDB^ by 0.011 in Fmax for BP. It could be attributed to the fact that the PDB training set for BP consisted of only 11,295 proteins, limiting the model’s ability to learn sufficiently. However, the comparison between GOBoost^All^ and GOBoost^All(PDB)^ revealed that the predictive performance of GOBoost^All^ improved significantly as the amount of trainable protein data increased.

### Performance of GO terms with different IC (Information Content) values on the PDB test set

We divided the functions into three groups, shallow, normal, and specific, according to their IC values to evaluate the performance of GOBoost in predicting functions of different difficulty levels. The larger the IC value, the higher the specificity value of the function, the lower the frequency of occurrence in the data set, and the more difficult it is to be predicted by the model. The experimental results in Table 4 show that GOBoost and GOBoost^All^ outperformed the HEAL method in all three categories of GO terms. Notably, in the group with the most difficult to predict IC > 10, GOBoost and GOBoost^All^ achieved improvements in AUPR and F1 scores for the three BP, CC, and MF sub-functional ontologies (52.48%, 16.96%), (31.09%, 12.94%), (13.28%, 9.68%), and (34.71%, 4.09%), (21.76%, 4.85%), (7.56%, 4.22%), respectively. While GOBoost achieved more significant improvement than GOBoost^All^, likely due to the influence of ensemble strategy, GOBoost^All^ also achieved considerable improvement in most evaluation groups, especially in the AUPR of IC *>* 10 group, which increased by 34.71%, 21.76%, and 7.56%. The improvement of GOBoost^All^ in Fmax for the group with IC *>* 10 is slightly inferior to other metrics, possibly due to the limited adjustment of the long-tail GO term weights, which aimed to optimize the overall performance of the model in this work. The integration of GOBoost^Tail^ effectively addresses this limitation, with GOBoost enhancing the performance over GOBoost^All^ by 0.044, 0.025, and 0.022 respectively.

**Table 4.** Performance of GO terms with different IC values on the PDB test set.

| GO | Method | F1 (↑) | | | AUPR (↑) | | |
| --- | --- | --- | --- | --- | --- | --- | --- |
|  |  | Shallow | Normal | Specific | Shallow | Normal | Specific |
| BP | GOBoost: | **0.732** | **0.586** | **0.400** | **0.808** | **0.547** | **0.369** |
|  | GOBoost<sup>All</sup>: | 0.715 | 0.556 | 0.356 | 0.788 | 0.505 | 0.326 |
|  | HEAL: | 0.686 | 0.525 | 0.342 | 0.750 | 0.428 | 0.242 |
| CC | GOBoost: | **0.409** | **0.665** | **0.349** | **0.833** | **0.644** | **0.506** |
|  | GOBoost<sup>All</sup>: | **0.409** | 0.659 | 0.324 | 0.812 | 0.627 | 0.470 |
|  | HEAL: | 0.336 | 0.620 | 0.309 | 0.779 | 0.510 | 0.386 |
| MF | GOBoost: | **0.481** | **0.761** | **0.442** | **0.935** | **0.815** | **0.674** |
|  | GOBoost<sup>All</sup>: | 0.471 | 0.738 | 0.420 | 0.932 | 0.783 | 0.640 |
|  | HEAL: | 0.464 | 0.697 | 0.403 | 0.917 | 0.739 | 0.595 |

Note: Shallow represents the protein functional group with $IC < 5$ , Normal represents the protein functional group with $5 \leq IC < 10$ , and Specific represents the protein functional group with $IC \geq 10$ . Data in bold indicates the best result.

### P-value study on PDB test set

We performed a P-value analysis to quantify the difference in prediction results between the GOBoost and HEAL methods. The P-values for MF, BP, and CC categories were of 8.38e-16, 3.67-e27, and 5.55e-08, respectively. There are significant differences in the prediction results of the HEAL method and GOBoost in the three types of functions. In particular, the prediction difference is the largest in the BP function, which is consistent with the previous experimental results. GOBoost has the most significant improvement in the BP function. In order to deeply explore the performance improvement of GOBoost compared with HEAL on protein graphs of different sizes, we divided proteins into five groups according to the number of amino acids in the protein graph. Then, we counted the distribution of the prediction differences between GOBoost and HEAL in each group. It can be seen from Fig. 3 that in any interval, the AUPR score difference between GOBoost and HEAL is mostly concentrated in the range of [-0.1, 0.2]. However, it is worth noting that the median of these differences always remains at and above 0, and the interval length between the upper quartile and the median is significantly longer than the lower quartile and the interval length between the median. This phenomenon shows that even for proteins whose prediction results have not been improved, the difference between GOBoost and HEAL is slight. In contrast, for those proteins whose prediction results have been improved, the improvement effect of GOBoost is more significant. The average improvement of each group shows that the improvement effect of GOBoost is not affected by the size of the protein graph. GOBoost has the best average improvement effect on BP and CC, even in the largest protein graph group. On MF, GOBoost has an average improvement of 0.074 in the 600-800 residue range and an average of 0.047 on the largest protein graph group, without too many extreme negative values.

**Fig. 3.**
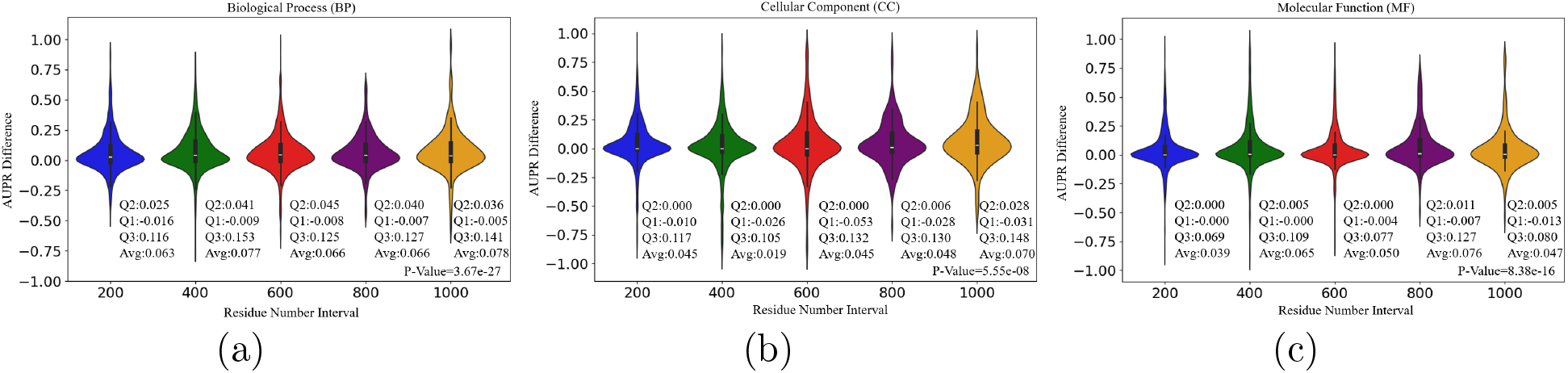
The P-value is calculated based on the AUPR values of all proteins predicted by GOBoost and HEAL. The x-axis is the number of protein residues, with each interval spanning 200, ranging from 0 to 1000. The y-axis is the difference between the AUPR values predicted by GOBoost and HEAL between [-1,1]. “Q2” is the median of each group of differences, “Q1” is the lower quartile, “Q3” is the upper quartile, and “Avg” is the mean of each group of differences. **(a)** Difference statistics on BP. **(b)** Difference statistics on CC. **(c)** Difference statistics on MF.

### Ablation study

#### Ablation study of long-tail optimization ensemble strategy

The ablation experiments for the long-tail optimization ensemble strategy are shown in Supplementary Table S1-2. This work first verified whether the integrated Head and Tail items are effective. The performance of both GOBoost^-Head^ and GOBoost^-Tail^ decreased, with GOBoost^-Tail^ experiencing a more significant decline than GOBoost^-Head^. This can be attributed to the fact that the distribution of GO terms of the three sub-functional ontologies is extremely unbalanced. Specifically, the number of GO terms belonging to the head distribution is smaller, which naturally results in a smaller impact on the model. Finally, from the comparison between GOBoost and GOBoost^All^, it can be seen that the entire optimization ensemble strategy proposed in this work significantly improved the predictive performance of the model in all dimensions. These highlight the importance of optimizing the ensemble model from a long-tail perspective.

Next, we analyzed the effects of different ensemble strategies. A key question is whether simply training the base models, GOBoost^Head^ and GOBoost^Tail^, separately and then connecting their predicted results (GOBoost^Ct^) would outperform training on all GO terms at once for GOBoost^All^. As shown in Supplementary Table S1-2, GOBoost^Ct^ significantly decreased on all evaluation metrics in both test sets. One important reason may be that both GOBoost^Head^ and GOBoost^Tail^ tend to focus more on the GO terms of the Head and Tail parts, which results in GOBoost^Ct^ being unable to capture the co-occurrence relationship between the complete GO terms. Moreover, the predicted distributions between the two base models may also differ, and directly connecting all predicted probabilities will lead to confusion in the final predicted probability distribution, further reducing the model’s performance. Finally, this work validated the impact of AVERAGE and Max optimization ensemble strategies. The experimental results show that the AVERAGE ensemble strategy achieves better results than the Max ensemble strategy. This may be because the Max strategy amplifies the erroneous GO terms predictions. On the other hand, the AVERAGE strategy averages the prediction results of the three base models, reducing the losses caused by extreme prediction errors and making the prediction results more stable.

#### Ablation study of long-tail design in the base model

We analyzed the effect of the long-tail design in the base model. Supplementary Table S3-4 provides detailed records of the ablation experiment results. Comparing the base model with the local label graph removed, GOBoost^All^ achieved the best performance in 10/15 evaluation metrics across two test sets. It is worth noting that the model’s predictability dropped most significantly on BP after removing the local label graph. The BP class functions are more complex and diverse, and the global label graph makes it difficult to capture such complex co-occurrence relationships completely. In contrast, the local label graph effectively supplements this. Compared with the base model with the long-tail term *γ*_*ht*_ removed from the loss function, GOBoost^All^ achieved the best results in 10/15 evaluation metrics. Finally, when both long-tail optimization designs were removed, the model showed a decline in 13/15 evaluation metrics. The long-tail design does effectively improve the predictive performance of the model.

Subsequently, this work validated the specific roles of global-local label graph (Classifier 2), average pooling (Classifier 1), and ℒ_*mgfl*_ loss in the model. It can be seen from Supplementary Table S3-4 that removing the complete global local label graph module resulted in a comprehensive decline in the three sub-functional ontologies. When evaluating BP function in the PDB test set, AUPR and Fmax decreased by 16.87% and 4.71%, respectively, and Smin increased by 5.94%. Compared to the variant base model without the average pooling operation, GOBoost^All^ achieved the best results in 13/15 evaluation metrics. Therefore, both the co-occurrence information between labels and the global information of the protein itself are essential. Combining the prediction results from these two directions can provide a more comprehensive prediction performance. Finally, compared to traditional binary cross-entropy loss, the ℒ_*mgfl*_ loss improved most evaluation metrics effectively, showing the rationality of the ℒ_*mgfl*_ loss design.

## Conclusion

In this work, we proposed a new method GOBoost to address the long-tail distribution effect of functional labels. We proposed a long-tail optimization ensemble strategy. Different from the traditional integration of deep learning methods and non-deep learning methods, it integrates three base models with different training objectives. Furthermore, we designed a global-local label graph module in the base model from two different perspectives, the global function, and the local individual function, to supplement the connection between long-tail labels and other labels. In addition, the ℒ_*mgfl*_ loss is designed to appropriately increase the weight of the long-tail GO terms, so that the model pays more attention to the long-tail function, thereby improving the overall prediction performance. We evaluated the performance of GOBoost on two datasets and compared it to the SOTA method; it achieved comprehensive improvements in all evaluation metrics on three sub-functional ontologies. In the more diverse PDB test set, obtaining the best results demonstrates that GOBoost can accurately predict the function of proteins with various features. GOBoost achieves the best results on the difficult AF2 test set, showing GOBoost’s capability to generalize and predict challenging proteins.

Designing models from the long-tail perspective has demonstrated great potential in protein function prediction. Looking ahead, we aim to leverage AlphaFold3 to enhance the accuracy of protein structure predictions for model training. As many protein functions are carried out through the interaction of multiple proteins, incorporating protein interaction network data could provide valuable contextual information, helping infer the biological processes and functions in which these proteins may participate. Additionally, we plan to integrate various non-deep learning methods, leveraging the strengths of each approach to select the most accurate predictions from a range of preliminary results.

## Key points

- Based on the long-tail distribution phenomenon of GO terms, we proposed and designed GOBoost, which achieved the best performance on all evaluation metrics of the test set.
- GOBoost constructed a global local co-occurrence label graph, which captured high-frequency functional labels and better supplemented the co-occurrence relationship between low-frequency labels and other labels.
- GOBoost designed loss functions at two granularities, namely individual positive and negative label imbalance and overall label distribution imbalance, to alleviate these two problems.
- GOBoost proposed an optimized integration strategy to alleviate the imbalance of functional labels.

## Supporting information

Supplemental file of GOBoost

## Author contributions statement

L. Z., Y. W and R. C. conceived the experiment(s), Y.W. and R.C. conducted the experiment(s), Y.W., L.Z., X.C., J.H., D.S. and R.C. analyzed the results. Y.W., L.Z., X.C., R.C., J.H., R.D. and D.S. wrote and reviewed the manuscript.

## Acknowledgments

This work was supported by the National Natural Science Foundation of China (No. 61976001), the Natural Science Foundation of Anhui Province (No. 2408085MF152), the Key Projects of the University Excellent Talents Support Plan of Anhui Provincial Department of Education (No. gxyqZD2021089), and Natural Sciences Summer Undergraduate Research Program by Pacific Lutheran University.

## Notes

https://cs.plu.edu/~caora/materials/GOBoost_supplementary.pdf

### Competing Interest Statement

The authors have declared no competing interest.

