## Supplemental file of GOBoost for "GOBoost: Leveraging Long-Tail Gene Ontology Terms for Accurate Protein Function Prediction"

### Supplementary Materials for GOBoost

#### 1. NUMBERING ITEMS IN THE SUPPLEMENTARY DOCUMENT

This file contains the following information:

Ablation study of long-tail optimization ensemble strategy on the PDB and AF2 dataSet.

Ablation study of long-tail design in the base model on the PDB and AF2 dataset.

#### 2. ABLATION STUDY OF LONG-TAIL OPTIMIZATION ENSEMBLE STRATEGY ON THE PDB AND AF2 DATASET.

Table S1: Ablation Study of long-tail optimization ensemble strategy on the PDB DataSet. Table S2: Ablation Study of long-tail optimization ensemble strategy on the AF2 DataSet.

#### 3. ABLATION STUDY OF LONG-TAIL DESIGN IN THE BASE MODEL ON THE PDB AND AF2 DATASET.

Table S3: .Ablation study of strategies based on the base model ($\mathrm{GOBoost}^{\mathrm{All}}$) using the PDB dataset. Table S4: .Ablation study of strategies based on the base model ($\mathrm{GOBoost}^{\mathrm{All}}$) using the AF2 dataset.

**Table S1.** Ablation Study of ensemble strategies on the PDB DataSet.

| Method | AUPR(↑) | | | Fmax (↑) | | | Smin(↓) | | |
|---|---|---|---|---|---|---|---|---|---|
| | MF | BP | CC | MF | BP | CC | MF | BP | CC |
| GOBoost | 0.765 | 0.458 | 0.573 | 0.787 | 0.659 | 0.745 | 0.292 | 0.450 | 0.401 |
| $GOBoost^{-Head}$ | 0.761 | 0.453 | 0.570 | 0.777 | 0.646 | 0.741 | 0.298 | 0.460 | 0.404 |
| $GOBoost^{-Tail}$ | 0.736 | 0.419 | 0.548 | 0.774 | 0.644 | 0.741 | 0.306 | 0.467 | 0.405 |
| $GOBoost^{All}$ | 0.733 | 0.415 | 0.546 | 0.769 | 0.637 | 0.739 | 0.309 | 0.471 | 0.408 |
| $GOBoost^{Ct}$ | 0.738 | 0.425 | 0.536 | 0.764 | 0.643 | 0.713 | 0.315 | 0.466 | 0.433 |
| $GOBoost^{Max}$ | 0.751 | 0.439 | 0.557 | 0.777 | 0.648 | 0.738 | 0.300 | 0.461 | 0.412 |

Note: To verify whether the long-tail optimization ensemble strategy is effective, the five variants of GOBoost are: (i) $GOBoost^{-Head}$: It represents the removal of $GOBoost^{Head}$ from GOBoost. (ii) $GOBoost^{-Tail}$: It represents the removal of $GOBoost^{Tail}$ from GOBoost.(iii) $GOBoost^{All}$: It indicates that the long-tail optimization ensemble strategy is not adopted. (iv) $GOBoost^{Ct}$: It represents the prediction result of directly connecting $GOBoost^{Head}$ and $GOBoost^{Tail}$. (v) $GOBoost^{Max}$: It integrates the highest prediction scores from three base models.

**Table S2.** Ablation Study of ensemble strategies on the AF2 DataSet.

| Method | MF | | BP | | CC | |
|---|---|---|---|---|---|---|
| | AUPR(↑) | Fmax (↑) | AUPR(↑) | Fmax (↑) | AUPR(↑) | Fmax (↑) |
| GOBoost | 0.582 | 0.556 | 0.246 | 0.497 | 0.318 | 0.643 |
| $GOBoost^{-Head}$ | 0.573 | 0.534 | 0.243 | 0.487 | 0.314 | 0.628 |
| $GOBoost^{-Tail}$ | 0.548 | 0.533 | 0.222 | 0.485 | 0.318 | 0.632 |
| $GOBoost^{All}$ | 0.540 | 0.526 | 0.220 | 0.480 | 0.313 | 0.621 |
| $GOBoost^{Ct}$ | 0.558 | 0.528 | 0.229 | 0.485 | 0.297 | 0.618 |
| $GOBoost^{Max}$ | 0.576 | 0.537 | 0.240 | 0.492 | 0.317 | 0.626 |

Note: To verify whether the long-tail optimization ensemble strategy is effective, the five variants of GOBoost are: (i) $GOBoost^{-Head}$: It represents the removal of $GOBoost^{Head}$ from GOBoost. (ii) $GOBoost^{-Tail}$: It represents the removal of $GOBoost^{Tail}$ from GOBoost.(iii) $GOBoost^{All}$: It indicates that the long-tail optimization ensemble strategy is not adopted. (iv) $GOBoost^{Ct}$: It represents the prediction result of directly connecting $GOBoost^{Head}$ and $GOBoost^{Tail}$. (v) $GOBoost^{Max}$: It integrates the highest prediction scores from three base models.

**Table S3.** Ablation study of strategies based on base model (GOBoost$^{\text{All}}$) using the PDB dataset.

| Method | AUPR(↑) | | | Fmax (↑) | | | Smin(↓) | | |
|---|---|---|---|---|---|---|---|---|---|
| | MF | BP | CC | MF | BP | CC | MF | BP | CC |
| GOBoost$^{\text{All}}$ | 0.733 | 0.415 | 0.546 | 0.769 | 0.637 | 0.739 | 0.309 | 0.471 | 0.408 |
| w/o L-label graph | 0.735 | 0.377 | 0.548 | 0.764 | 0.602 | 0.733 | 0.312 | 0.499 | 0.415 |
| w/o L-Loss | 0.725 | 0.418 | 0.533 | 0.756 | 0.639 | 0.733 | 0.318 | 0.469 | 0.412 |
| w/o L-both | 0.734 | 0.374 | 0.521 | 0.758 | 0.601 | 0.715 | 0.318 | 0.501 | 0.436 |
| w/o GL-label graph | 0.703 | 0.345 | 0.526 | 0.744 | 0.607 | 0.734 | 0.337 | 0.499 | 0.413 |
| w/o Mean-P | 0.718 | 0.400 | 0.518 | 0.761 | 0.630 | 0.730 | 0.318 | 0.479 | 0.421 |
| w/o $\mathcal{L}_{mgfl}$ | 0.739 | 0.400 | 0.547 | 0.767 | 0.632 | 0.735 | 0.318 | 0.475 | 0.410 |

Note: To verify the effectiveness of the long-tail design in the base model, the six variants of GOBoost$^{\text{All}}$ are: (i) w/o L-label: This variant removes the local label graph from the global local label graph module. (ii) w/o L-Loss: This variant removes the long-tail term of the $\mathcal{L}_{mgfl}$ loss. (iii) w/o L-both: This variant also removes the local label graph submodule in the global-local label graph module and the long-tail term of the $\mathcal{L}_{mgfl}$ loss. (iv) w/o GL-label graph: This variant removes the entire global local label graph module, and the base model only has one prediction head after average pooling. (v) w/o Mean-P: This variant removes the average pooling prediction head and only retains the prediction head after the global local label graph module. (vi) w/o $\mathcal{L}_{mgfl}$: This variant replaces the $\mathcal{L}_{mgfl}$ loss with a binary cross entropy loss.

**Table S4.** Ablation study of strategies based on base model (GOBoost$^{\text{All}}$) using the AF2 dataset.

| Method | MF | | BP | | CC | |
|---|---|---|---|---|---|---|
| | AUPR(↑) | Fmax (↑) | AUPR(↑) | Fmax (↑) | AUPR(↑) | Fmax (↑) |
| GOBoost$^{\text{All}}$ | 0.540 | 0.526 | 0.220 | 0.480 | 0.313 | 0.631 |
| w/o L-label graph | 0.543 | 0.519 | 0.220 | 0.465 | 0.322 | 0.627 |
| w/o L-Loss | 0.547 | 0.513 | 0.221 | 0.480 | 0.307 | 0.615 |
| w/o L-both | 0.551 | 0.520 | 0.218 | 0.478 | 0.302 | 0.624 |
| w/o GL-label graph | 0.535 | 0.516 | 0.210 | 0.478 | 0.300 | 0.619 |
| w/o Mean-P | 0.523 | 0.506 | 0.222 | 0.489 | 0.311 | 0.611 |
| w/o $\mathcal{L}_{mgfl}$ | 0.530 | 0.521 | 0.229 | 0.485 | 0.304 | 0.626 |

Note: To verify the effectiveness of the long-tail design in the base model, the six variants of GOBoost$^{\text{All}}$ are: (i) w/o L-label: This variant removes the local label graph from the global local label graph module. (ii) w/o L-Loss: This variant removes the long-tail term of the $\mathcal{L}_{mgfl}$ loss. (iii) w/o L-both: This variant also removes the local label graph submodule in the global-local label graph module and the long-tail term of the $\mathcal{L}_{mgfl}$ loss. (iv) w/o GL-label graph: This variant removes the entire global local label graph module, and the base model only has one prediction head after average pooling. (v) w/o Mean-P: This variant removes the average pooling prediction head and only retains the prediction head after the global local label graph module. (vi) w/o $\mathcal{L}_{mgfl}$: This variant replaces the $\mathcal{L}_{mgfl}$ loss with a binary cross entropy loss.
